# Children’s DNA Methylation and Family Dynamics in a Congo Basin Subsistence Community: Links with Parental Conflict and Fathers’ Caregiving

**DOI:** 10.64898/2026.06.15.732362

**Authors:** Meingold Hiu-ming Chan, Sarah M. Merrill, Beryl Zhuang, David T.S. Lin, Julia L. MacIsaac, Valchy Miegakanda, Sheina Lew-Levy, Adam H. Boyette, Michael S. Kobor, Lee T. Gettler

## Abstract

Family environments may contribute to children’s long-term health through biological processes, including epigenetic regulation such as DNA methylation (DNAm). However, most studies in this area focus on Euro-American populations while also rarely including fathering data. The current study investigated children’s blood DNAm associations with positive (father caregiving) and negative (parental conflict) family dynamics in a smaller-scale subsistence society living in the Congo Basin rainforest. We measured DNAm from dried blood spots of 54 children (mean age=8.48 years) and conducted three epigenome-wide association studies aimed at discovering differential co-methylated regions (CMRs) associated with family dynamics. Via path models, we investigated the health implications and shared contribution of family factors of the identified CMRs. Differential DNAm associated with family dynamics was localized to genes related to stress, immunology, development, and aging, thus possibly linking to children’s physical health and were simultaneously connected to other family factors such as number of siblings. Our findings suggested similarities in biological embedding of family factors across socio-ecologically diverse contexts.

## Introduction

Family context, including both interparental and parent-child dynamics, has a profound impact on child health and development. Developmental models such the Developmental Origins of Health and Disease (DOHaD) framework and the Biological Embedding of Early Experience, posit that early life exposures shape trajectories of biological development with implications for long-term health and aging (Barrett, 2017; Hertzman, 1999). Early life adversity within family systems, which includes a range of exposures such as maltreatment, negative parenting behaviors, and parental conflict, has been widely studied and found to affect children’s physical growth and health, mental well-being, and numerous other developmental (Harold & Sellers, 2018; Wade et al., 2022). Indeed, stressful family experiences can take a biological toll on children. Dysregulated stress physiology, inflammatory (e.g., CRP, IL-1/6/10, TNF-α), metabolic, and cardiovascular indicators have all been documented as potential pathways through which family dynamics affects children’s functioning (Boyce & Kobor, 2015; Chiang et al., 2022; Gettler, Lin, et al., 2020; Harold & Sellers, 2018). Meanwhile, results from systematic reviews and meta-analyses indicate that positive family experiences, such as parental caregiving and warmth, attachment security, and safe and protective environments, can promote child health and development or buffer the biological impacts of negative experiences (Jeffries Hein et al., 2024; Kallapiran et al., 2025). For example, a recent review discussed consistent, negative association between positive parenting behaviors (i.e., parental responsiveness and warmth) with child pro-inflammatory markers (O’Brien et al., 2023). Building on these findings, the DOHaD framework has broadened its focus beyond early life adversity to also include positive childhood experiences — an area that remains comparatively understudied (Jeffries Hein et al., 2024). The more recently proposed Paternal Origins of Health and Disease (POHaD) further extends this framework to highlight the importance of including paternal influences in addition to the more common focus of maternal impact (Soubry, 2018), which parallels broader calls for more rigorous study of paternal effects in child development research. Our understanding of fathers’ contributions to children’s development through high-quality caregiving has also grown substantially and extended to children’s developmental physiology (Gettler & Barr, 2022).

### Biological embedding & DNAm associations of family dynamics in children

The myriads of biological associations found with children’s early family experience reviewed above lend supports to the biological embedding hypothesis. The underlying molecular pathways of this process have been increasingly studied in the past decade. Specifically, epigenetic changes—molecular modifications to the genome without changing the underlying DNA sequences (Aristizabal et al., 2020)—may offer further insight into how family dynamics becomes biological embedded in children (Boyce & Kobor, 2015). DNA methylation (DNAm) is an epigenetic mechanism that in part captures environmental exposures and varies systematically across the genome, making it a powerful tool for investigating how early-life experiences shape long-term biology. DNAm is a well-characterized, mitotically heritable epigenetic mark that can be stably maintained over time and leads to changes in biological functioning that has implication to human development and health (Aristizabal et al., 2020; Jones et al., 2018).

Although DNAm studies with candidate gene approach have provided preliminary evidence that family dynamics, including parental conflict and parental caregiving, can become biologically embedded through DNAm (Provenzi et al., 2020; Unternaehrer & Meinlschmidt, 2016), more comprehensive epigenome-wide association studies (EWAS) have now become the standard approach to investigate differential DNAm associations with psychosocial factors (Zhang & Liu, 2022). Several EWASs have linked differential DNAm to cumulative family stress (Bush et al., 2018; Chan et al., 2025; Essex et al., 2013), including marital conflict reported by mothers (Bush et al., 2018) and both parents (Essex et al., 2013), particularly in genes involved in stress response and immune function (Zhang & Liu, 2022). However, there remains a scarcity of EWASs on more positive aspects of family environment such as high quality parental caregiving (Provenzi et al., 2020). A construct strongly related to caregiving quality and can be impacted by parental conflict is attachment style, which was found to be linked to differential DNAm in recent EWASs, again in genes related to stress responses, inflammation, and cognitive development (Garg et al., 2018; Merrill, Gladish, et al., 2021). Supportive family environment and positive parenting has also been shown to alleviate the negative impact of psychosocial stress as reflected by biological aging measured with DNAm-derived epigenetic clocks (Brody et al., 2016; Sullivan et al., 2023).

The role of fathers in shaping children’s developmental biology remains understudied in humans, with the limited existing work focusing mainly on stress-responsive physiology and showing that effects vary by the socio-ecological context of fathers’ caregiving and family roles (Gettler, 2016; Gettler & Barr, 2022). Given the sparse literature and general underrepresentation of fathers in research, it is perhaps unsurprising that no EWAS on paternal caregiving has been reported to date. Nevertheless, the influence of fathers on offspring DNAm has been hypothesized (Ferber et al., 2021) and is beginning to receive empirical attention. Specifically, fathers’ adverse childhood experiences and psychopathological symptoms are associated with differential DNAm of their infant offspring (Gettler & Barr, 2022; Merrill, Moore, et al., 2021). Thus, this emerging body of research is supporting the idea of both positive and negative family dynamics, including the role of fathers, becoming biologically embedded at the molecular level and reflected through differential DNAm.

### Ethnographic context & the biological implication of parental conflict and father caregiving in Bandongo children

To shed light on the generalizability of the biological embedding of family dynamics, studying this process in diverse contexts is important, especially populations that are culturally, socio-ecologically, and economically different from the commonly studied Euro-American societies. The current study drew on data from Bandongo children living in a smaller-scale, subsistence fishing-farming society located in the northern Republic of the Congo. This community is located remotely in the rainforest far from any urban center, with no direct road access and relatively little market integration and characterized by high degrees of pathogen stress and highly active subsistence lifestyles, creating energetically demanding daily conditions (Gettler, Lin, et al., 2020). Moreover, the local cultural context differs substantially from industrialized and post-industrialized societies. Bandongo society is patriarchal, with fairly rigid social hierarchy based on gender, age, and both ascribed and acquired status. Thus, older men in this village generally have higher social status. In the family context, verbal and occasionally physical disputes between spouses are not uncommon, resulting in exposure to parental conflict among children (Boyette et al., 2018; Gettler, Lin, et al., 2020). However, adults from this community noted that “children cry and ‘hold it in their heart’” (Boyette et al., 2018, p. 841) when exposed to parental conflict and reported that a component of being a good father in Bandongo society is avoiding spousal conflict. Other paternal roles in Bandongo society are multidimensional. Fathers are highly valued for their indirect caregiving, particularly their provisioning of food and resources, and resource acquisition also confers social status in the community. Fathers are comparatively less involved with hands-on caregiving of young children. Bandongo children primarily receive direct care from their mothers and other family members, including older siblings and grandmothers (Boyette et al., 2020). Nonetheless, both paternal indirect and direct caregiving have been linked to aspects of child health and development, including energetic condition and stress physiology (Boyette et al., 2018, 2020). Ethnographic observation of this cultural group also suggested that the Bandongo kin networks are independent economic units, with families generally being responsible for their own production/consumption (Boyette et al., 2018, 2020). High fertility and large families are culturally valued, in part due to children’s contributions to subsistence and household labor, including childcare. However, the rainforest ecology is also pathogen-intensive and energetically-demanding, meaning more children also increases provisioning demands (Boyette et al., 2020; Gettler, Lin, et al., 2020). Taken together, Bandongo children’s health is thus likely affected by multiple family factors.

Biological embedding of parental conflict and father caregiving has been documented in the Bandongo societies with multiple biomarkers. For example, parental conflict predicted heightened stress physiological responses and elevated biomarkers of physiological stress, indicated by Epstein-Barr virus (EBV), in children (Boyette et al., 2018). Fathers’ direct and indirect care were linked to better physical health and growth in their children (Boyette et al., 2018), although indirect care is more culturally valued than direct care (Boyette et al., 2020). A prior analysis of this cohort’s DNAm data reported a significant association between parental conflict and DNAm-derived epigenetic age acceleration (EAA) (Gettler, Lin, et al., 2020).

Specifically, Bandongo children in families with higher levels of parental conflict were found to have greater intrinsic EAA—faster epigenetic aging, independent of differences in blood cell type proportions—compared to their counterparts. Although less is known about EAA-health associations in children than in adult populations, previous studies have linked greater EAA in children and adolescents with health outcomes, including growth, asthma, and allergy (Gettler, Lin, et al., 2020). Research from a multi-decade birth cohort study in urban Philippines has also shown that early life poverty, which could influence family dynamics, was linked to differential DNAm of genes involved in immune system and skeletal development (McDade et al., 2019). Therefore, despite the paucity of research in societies outside of Euro-American contexts, emerging biological findings suggest differential DNAm associated with family environment present even in societies socio-ecologically different from those typically studied.

### The Current study

The current study leveraged data collected from a sample of Bandongo children, allowing us to broaden the range of contexts in which we study the biological embedding of family dynamics. First, we adopted an EWAS approach to test for differential DNAm regions associated with positive (father caregiving) and negative (parental conflict) family dynamics, as rated by father peers within the Bandongo society. Given previous findings on health and biomarkers in this cohort, we hypothesized that identified DNAm regions would reside within genes associated with stress and growth (H1). To evaluate the health implications of identified DNAm regions, we used a multivariate approach to test our hypothesis that the identified DNAm regions will be associated with differences in children’s health (e.g., EBV, C-reactive protein; CRP) and energetic condition (e.g., weight-for-height; WFH, skinfold thickness; SFT) (H2). We further contextualize our findings by examining the shared contribution of both father caregiving and parental conflict on identified DNAm regions while considering the effect of other potentially relevant family factors on our main variables of interest in a path model. We hypothesized that father caregiving and parental conflict will have shared contribution on identified DNAm regions (H3) and explored whether fathers age and number of children in the household contribute to the associations.

## Methods

### Participants

The study sample was recruited from a Bandongo village in the rainforest of the Likouala Department, Republic of the Congo, and has been previously described (Boyette et al., 2018; Gettler et al., 2019; Gettler, Lin, et al., 2020). The village has a population of less than 200 individuals who subsist largely on fishing, farming, and hunting, with some small-scale, local commerce (Boyette et al., 2018). Households with at least one child under the age of 18 were eligible for this study, and all 20 eligible families in the village participated. Only children under age 18 were included in analyses in this study and the final sample was 54 children from 17 families (**Supplementary Table 1**). The study was approved by the village council through a public meeting and by the Institut de Recherche en Sciences Exacts et Naturelles in Brazzaville. Verbal consent from adult participants and assent from children were obtained individually.

Institutional Review Boards of Duke University (Protocol # 2017 0038) and the University of Notre Dame (# 18 02 4397) had approved the study.

### Measures

#### Children’s anthropometrics and biomarkers

Children’s weight in kilograms (kg) and height in centimeters (cm) were measured according to standardized measurement procedures (Lohman et al., 1988). WFH, an indicator for longer-term energetic condition, was computed, adjusted for age and sex, and then transformed to z-scores (Boyette et al., 2018; Gettler, Lin, et al., 2020). Triceps SFT, often assessed as an indicator of nutritional status, was measured using Lange calipers in millimeter (mm) and transformed to z-scores, after accounting for age and sex (Boyette et al., 2018).

As previously described in detail (see Boyette et al. and Gettler et al. 2021), blood by finger-prick sampling was stored as dried blood spots (DBS) on filter paper. DBS samples were assayed for EBV and CRP levels using enzyme-linked immunosorbent assay approaches at the Global Health Biomarker Laboratory at the University of Oregon (**Supplementary Method**).

#### Peer rankings for family dynamics and child health

Villagers were informally interviewed by the research team to understand locally-valued roles for fathers and family dynamics in the Bandongo community. These interviews focused on the responsibilities of a father, marital dynamics, and healthy child development. Informed by qualitative analyses on these interviews (Boyette et al., 2018), a ranking task was used to obtain peer perspectives on fathering quality and child well-being in four domains: indirect care (providing food/resources), direct care (childcare and rearing), parental conflict (conflict with wife/wives), and children’s health. In this task, fathers were asked questions on parental conflict (“*Who disputes with their wife/wives the most?*”) and paternal care, including two types of paternal care, direct care (“*Who ‘attends to’ their children the most*?” and “*who is most likely to stay home/sacrifice other activities when their children are sick to care for them*?”) and indirect care (“*Who works the hardest?*”). Peer-ranking tasks on core, shared cultural domains (e.g., local plant knowledge; hunting ability) are commonly used in studies of smaller-scale societies where community familiarity is high. The Bandongo men in the present sample knew each other well, as most had grown up together and/or were related through kinship by various degrees (Gettler, Lew-Levy, et al., 2020). For each of the peer-ranking questions, participants were asked to use a photo array of their peers to rank fathers from the rest of the participating households (excluding himself) from low (last position is coded as 1) to high, while ties were allowed. The average of the rankings was taken as the summary ranking score for each domain per father. Reliability statistics for the rankings showed appropriate consensus between fathers (Cronbach’s alpha: 0.84-0.95) (Boyette et al., 2018). Higher ranking for the conflict question indicated peer perception of more parental conflict and greater paternal care for the paternal care questions. Children from the same household received the same ranking scores in each of the domains.

### Biological sample processing

#### DNA extraction, DNA methylation arrays, and preprocessing

Genomic DNA was extracted from the DBS samples using a combined protocol involving GenTegra GenSolve Extraction Kits and Qiagen QIAamp DNA Mini Kits and bisulfite-converted using the Zymo EZ DNA Methylation Kits. Samples were randomized on the array against age, sex, and household to minimize technical confounds and 160 ng of bisulfite-converted DNA was applied to the Illumina Infinium MethylationEPIC Beadchips (EPIC v1) as per manufacturer’s protocols (Illumina Inc. USA); details were previously described (Gettler, Lin, et al., 2020). DNAm array data quality check, preprocessing, and downstream analysis were performed in R version 4.2.2. All 54 children’s DNAm array data passed quality checks, which included identifying poor quality samples by average detection p-values. Potential sample cross contamination was checked by hierarchical clustering analysis of the 59 SNP probes with no indication of sample contamination. SNP probes, probes mapped to sex chromosomes, and cross reactive probes and polymorphic probes (Pidsley et al., 2016) were removed. Furthermore, low quality probes that either had beadcount <3 in three or more samples, or with a detection p-value > 0.05 in any sample were filtered. 788,032 probes were carried forward to normalization. Raw data were background and color corrected in Illumina GenomeStudio Software (version 2.0, Illumina Inc.), and followed by Beta MIxture Quantile dilation (BMIQ) normalization (Teschendorff et al., 2013). Another 43,516 probes with missing values were excluded prior to batch correction. Row and chip batch effects were removed using the *ComBat* function in the *sva* R package (Leek et al., 2012). A final count of 744,516 probes was retained for subsequent analysis. DNAm β value per probe is ratio of the methylated probe intensity over the total of methylated and unmethylated probe intensities, with a range of 0-1 (Du et al., 2010).

#### Analyses Computational cell type proportion estimation

Cellular composition is one of the main contributors to DNAm profile variations (Jones et al., 2018; Zheng et al., 2017). Referenced-based methods to computationally estimate cell type proportions can serve as surrogates in lieu of cell counts (Houseman et al., 2012; Salas et al., 2018) (**Supplementary Figure 1**). Relative cell type proportions of each sample were predicted by the constrained projection method (Houseman et al., 2012), with IDOL reference library (Salas et al., 2018) by using the estimateCellCounts2 function in the R package FlowSorted.Blood.EPIC (version 2.2.0). Relative proportions of six cell types in blood: neutrophils, monocytes, B lymphocytes (Bcells), T helper lymphocytes (CD4T), T cytotoxic lymphocytes (CD8T), and natural killer lymphocytes (NK) were estimated. Such cell type estimations are positive and proportional, which sum up to 1; and the proportions of cell types are correlated. Cell type estimations were transformed by the isometric logratio (ilr) transformation for compositional data followed by robust principle component analysis (PCA) (Filzmoser et al., 2009). The top three principal components (PCs) accounted for 89.8% of predicted cell type proportions variability and were included as covariates in the statistical models to account for this.

#### Co-methylated Regions as Unit of Analysis

Because DNAm data involves hundreds of thousands of probes and our sample is necessarily small—having recruited all eligible participants from a single village—the stringent multiple testing corrections required may obscure genuine biological signals. Region-based analysis reduces redundant measurements and noise while capturing DNAm patterns that are consistent and more likely to be biologically relevant in the data (Gatev et al., 2020). Closely located cytosine-guanine (CpG) dinucleotides sites tend to have correlated methylation states and are more likely functioned as biological units (Gatev et al., 2020; Hui et al., 2018). Thus, in the current study, the unit of analysis employed was co-methylated regions (CMRs), which are defined as groups of adjacent CpGs with correlated DNAm levels across samples (Gatev et al., 2020). CMRs were constructed based on the 744,516 probes that passed quality check using the *cmr* function in the *CoMeBack* R package (version 1.0) (Gatev et al., 2020) (details in **Supplementary Method**). A final count of 45,907 variable CMRs was included in the association analysis (**Supplementary Figure 2**). A composite DNAm β value of a CMR (CMR β) was represented by the median DNAm β value of all CpGs within the CMR.

#### EWAS for family dynamics

Three EWAS using robust linear mixed-effects (lme) models including a random intercept for family (n = 17) was employed to discover associations between Bandongo community children’s (n = 54) CMR β value and the three fathers’ peer rated family dynamics variables (parental conflict, direct and indirect paternal care) (Koller, 2016). The denominator degrees of freedom were estimated from the fitted lme model. Because multiple children were sampled from the same households, and the key predictors (paternal caregiving and parental conflict rankings) were measured at the family level, all models included a random intercept for family. This multilevel specification allowed family-level predictors to be evaluated in relation to child-level outcomes while appropriately adjusting standard errors for within-family clustering (Koller, 2016). Age, sex, and the top three estimated cell type proportion PCs were included as covariates, as these variables are known to contribute to DNAm levels (Jones et al., 2018).

Accordingly, inference for key predictors was constrained by the number of family-level exposure units, whereas child-level covariates were estimated using child-level variation. The robust lme model was applied to each of the 45,907 variable CMRs:

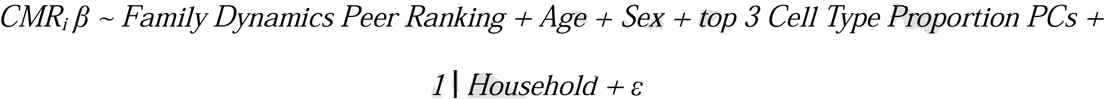

Models were run using the *rlmer* function implemented in *robustlmm* R package (version 3.1.2) with default settings. Family dynamics ranking β coefficient estimates and *t* statistics were extracted from each model. *P*-value was calculated from *t* value using Satterthwaite approximations of degrees of freedom (Geniole et al., 2019). Benjamini-Hochberg procedure was then applied across 45,907 tests to calculate adjusted *p*-values for each key predictor (Benjamini & Hochberg, 1995), with false discovery rate set at 0.2 (Korthauer et al., 2019). Post-correction, CMRs with an adjusted *p*-value <0.2 were determined as medium-confidence associations and adjusted *p*-value <0.05 were high-confidence associations with peer ranking family dynamics. A technical effect size threshold was set at |CMR Δβ| > 0.03 for the 95^th^ and 5^th^ percentile range of each predictor based on literature recommendations and similar analyses in pediatric blood (Merrill, Gladish, et al., 2021; Merrill, Moore, et al., 2021) to increase confidence in the changes observed being greater than technical noise. This technical threshold allows us to test whether the estimated DNAm difference across a high-versus-low peer-ranking contrast is larger than the average technical difference observed when the same individual is assayed twice. We computed CMR Δβ as:

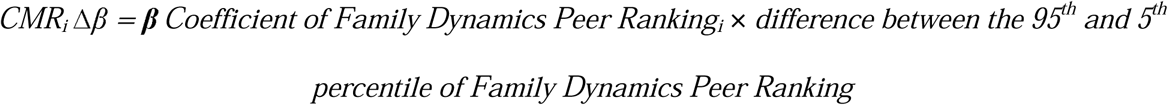

β coefficient of family dynamics peer ranking for each CMR_i_ Δβ was obtained from the exploratory robust lme model.

#### DNA methylation quantitative trait loci (mQTL) analysis

DNA methylation levels of CpGs can be affected by nearby single nucleotide polymorphisms (SNPs), termed methylation quantitative trait loci (mQTLs) (Bell et al., 2011; Smith et al., 2014). Thus, we examined whether any identified CMRs were potentially associated with mQTLs. As the analyses did not have available genotype data from our cohort and available mQTL databases were inappropriate for this population (Shang et al., 2023), we inspected the modality of the CMR β value distribution since CpGs associated with mQTLs often have bi- or trimodal distributions, reflecting DNAm differences in multiple genotypes (Schröder et al., 2017; Smith et al., 2014). We employed the *nmode* function from R package *ENmix* (version 1.34) to identify CMR β values with multi-modal distribution using default parameters (Xu et al., 2016).

#### Path models

Path analysis is well-suited for multivariate analyses since it allows for simultaneous estimations of associations among multiple variables, including more than one dependent variable, while accounting for covariance among predictors. Two path models were run to characterize the identified family dynamics-associated CMRs. The first model examined the associations with child health outcomes to evaluate the relevance of identified CMRs in the context of child health. The second model then examined the shared contribution of all family-dynamics measure simultaneously on the identified CMRs (**Supplementary Figure 5**). All path analyses were run with R *lavaan* package (Rosseel, 2012). Path coefficients were estimated using maximum likelihood (ML). There were no missing data on any of the variables included in the path analyses. We evaluated the goodness of fit of our models using comparative fit index (CFI), the root mean square error of approximation (RMSEA), and standardized root mean square residuals (SRMR) in which a model meeting the following criteria was considered a good fit: CFI > .95, RMSEA < .06, and SRMR < .08, while models with CFI of .90–.95 and RMSEA < .08 was considered acceptable fit (Hu & Bentler, 1995). Path estimates were all standardized. Standardized estimate ranged from |0.10-0.30| is considered as a small effect size, ranged from |.30-.50| as a moderate effect size, and above |.50| as a large effect size (Cohen, 2016).

Because our sample size did not support multivariable modeling of all identified regions directly, the path analyses used dimension-reduced summaries of the identified CMRs of PCA with Promax rotation on the identified CMRs from each EWAS. Only CMRs that had a loading larger than |0.4| was included for parsimony and to facilitate interpretation (Stevens, 2002).

Three sets of CMR PCs that were parental conflict-associated, direct care-associated, and indirect care associated, respectively, were used in the path models. These PCs should be interpreted as composite DNAm patterns associated with each family domain, rather than as isolated locus-specific effects. Furthermore, the loadings of the CMR should be considered in conjunction with its initial directionality of EWAS associations for interpretation of downstream analyses. For example, when the PC loadings and the EWAS associations of the CMRs shared the same directionality, the constructed PC should be expected to have a positive association with the respective variable of interest. In contrast, when the PC loadings (e.g., positive) and the EWAS associations of the CMRs (e.g., negative) had opposite directionality, the constructed PC should be expected to have a negative association with the respective variable of interest. A PC with mixed directionality (some loadings have consistent directionality with the EWAS associations while some do not) might be less interpretable.

In the first path model, the CMR PCs were the predictors, whereas the outcomes were five child health measures. This included biomarkers of inflammation and psychosocial stress (CRP and EBV, respectively), physical growth (WFH and SFT), as well as peer-ranked child health. We adjusted for the top three cell type PCs and child age in the model since they are known to drive DNAm variations and allowed all CMRs to covary with each other and all child health outcomes to covary with one another.

In the second path model, the CMR PCs were included as outcome variables with the three family dynamics variables as predictors. Additionally, we included father’s age and total number of children in the model given their relevance to family dynamics in the Bandongo cultural context (Gettler et al., 2019). Specifically, we estimated the paths in which father’s age predicted the total number of children and the peer-ranking family dynamics variables. We also estimated the paths from total number of children to family dynamics variables and CMR PCs (**Supplementary Figure 5)**. Similarly, we adjusted for the top three cell type PCs and child age in the model and allowed all CMRs to covary with each other and all family dynamics variables to covary with one another. Insignificant paths were removed in the final model for parsimony.

## Results

Using data from families in the Bandongo fishing-farming society, our study investigated the biological embedding of family dynamics in this cultural context through the lens of epigenetics (**Figure 1**). Mean age of children and fathers was 8.48 and 38.29, respectively, and about half of the children were female. On average, each household had around six children, with a range of two to 15 children (**Supplementary Table 1**). Pearson’s correlations showed that the peer-ranked variables were not significantly correlated with each other (*r*s = −0.09 – 0.02, *p*s > 0.05) (**Supplementary Figure 3**).

**Figure 1.**
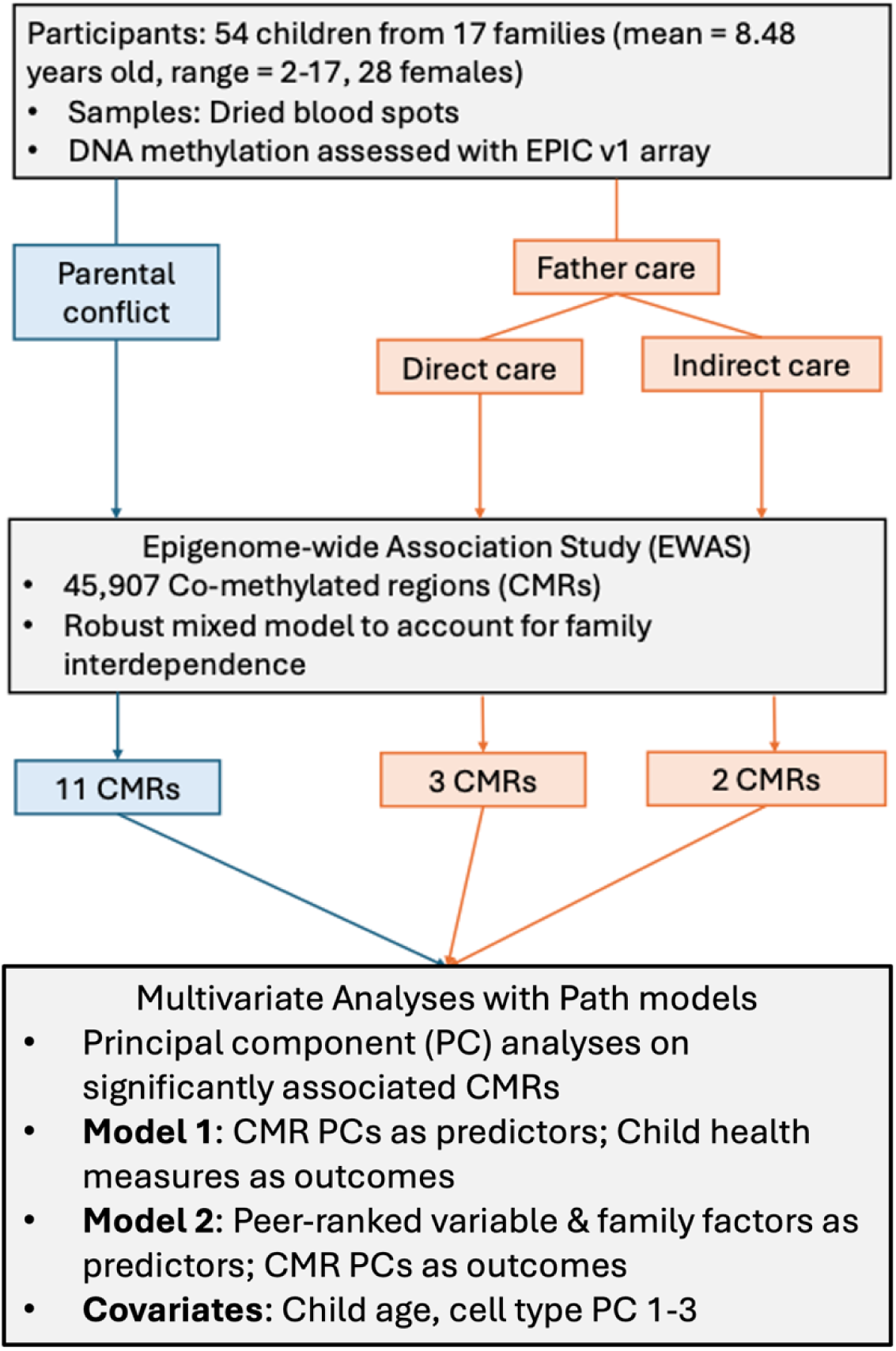
Study Overview. The first aim of the study was to discover differential DNA methylation of co-methylated regions (CMRs) associated with positive (father caregiving) and negative (parental conflict) family dynamics, using epigenome-wide association study. The second aim was to evaluate the health implications and shared contribution of family factors of identified CMRs using a multivariate approach.

### CMRs associated with parental conflict (one high-, ten medium-confidence) and paternal care (five medium-confidence)

Bandongo children’s genome-wide DNAm associations with peer-ranked family dynamics were explored with a region-based approach, which are defined as groups of adjacent CpGs with correlated DNAm levels across samples (Gatev et al., 2020). With 45,907 variable CMRs tested, one high-confidence CMR (adjusted *p* < 0.05, *p* = 3.04E-07, Δβ = 0.037) and ten medium-confidence CMRs (adjusted *p* <0.20, *p* < 4.87E-05,) were identified to be associated with parental conflict (**Figure 2**; **Table 1**). Five CMRs, all medium-confidence, were found to be linked to peer-ranking for paternal direct and indirect care (**Figure 3**; **Table 1**). Two of the conflict-associated CMR and one indirect care-associated CMR showed bimodal β distribution, suggesting genetic control (**Table 1; Supplementary Figure 4**).

**Figure 2.**
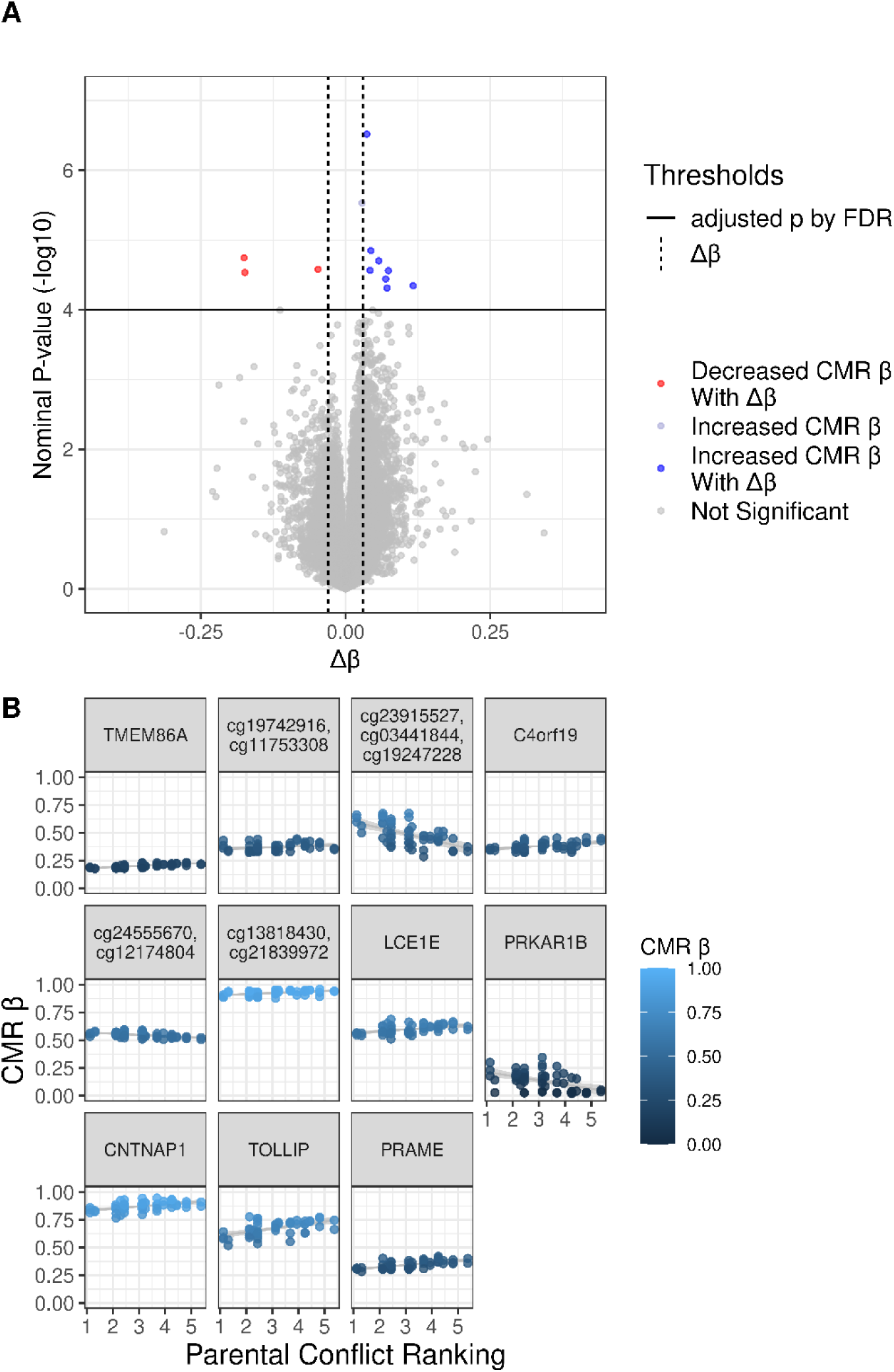
DNA methylation at 11 co-methylated regions (CMR) significantly associated with parental conflicts. Results of the associations between median DNAm level of each co-methylated regions (CMR) and parental conflict were tested by robust linear mixed-effects model while controlling for child’s sex, age, and estimated cell-type proportions, with a random intercept for family. (a) Volcano plot showing that eleven CMRs passed the statistical (adjusted p-value < 0.2) and technical thresholds (|Δβ |> 0.03), represented by the dashed line. (b) Line graphs showing the relation between parental conflict on the x-axis and β value ranged from 0 to 1 on the y-axis for the 11 CMRs. The colors of the data points indicate β values. Trend lines and confidence intervals are shown.

**Figure 3.**
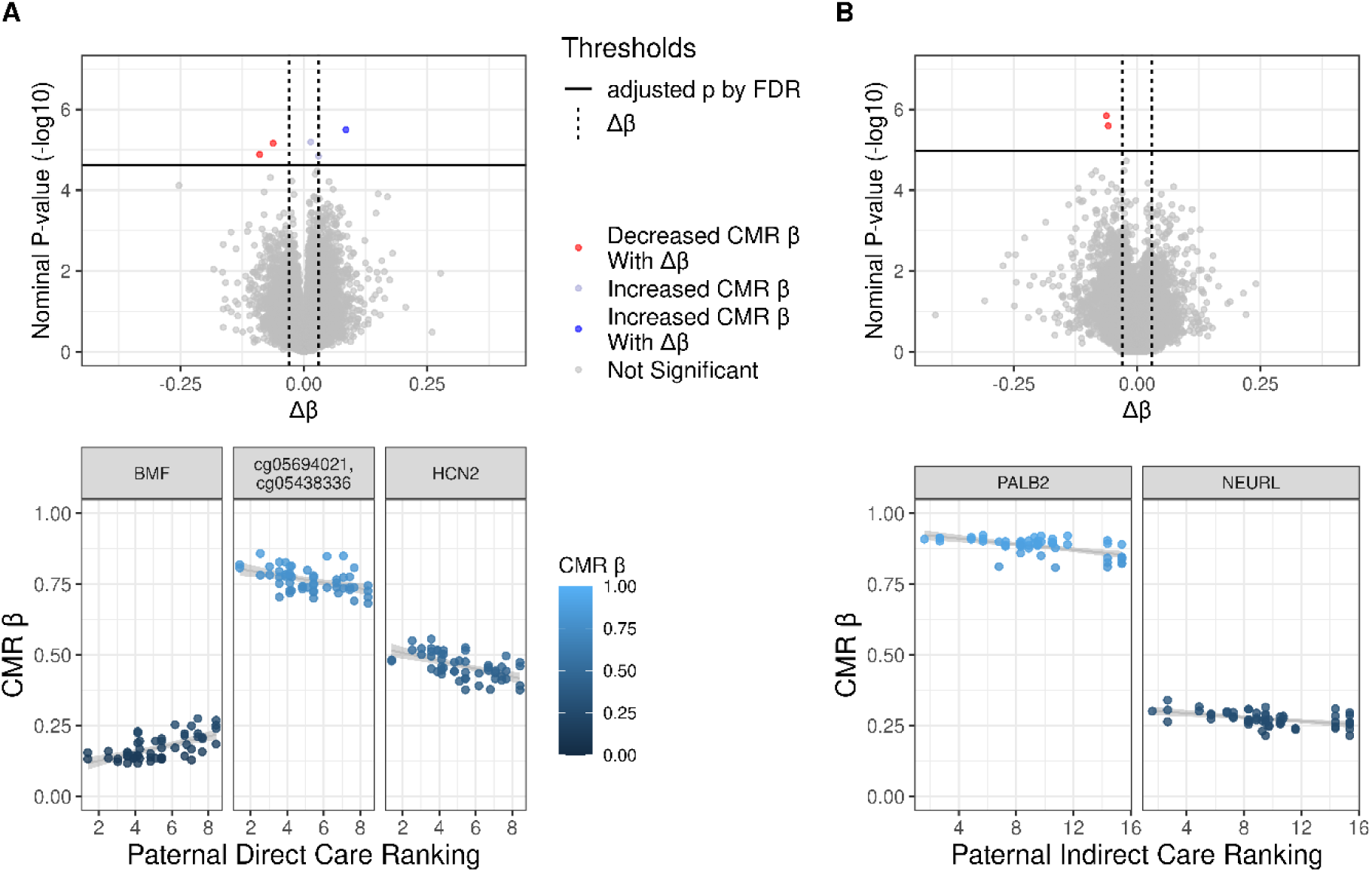
DNA methylation at 5 co-methylated regions (CMR) significantly associated with paternal conflicts. Results of the associations between median DNAm level of each co-methylated regions (CMR) and paternal direct and indirect care were tested by robust linear mixed-effects model while controlling for child’s sex, age, and estimated cell-type proportions, with a random intercept for family. (a) Volcano plot showing that three and two CMRs for direct and indirect care, respectively, passed the statistical (adjusted p-value < 0.2) and technical thresholds (|Δβ |> 0.03), represented by the dashed line. (b) Line graphs showing the relation between paternal care on the x-axis and β value ranged from 0 to 1 on the y-axis for the 5 CMRs. The colors of the data points indicate β values. Trend lines and confidence intervals are shown.

**Table 1.** CMRs significantly associated with parental conflict and paternal care.

| Variables | CMR CpGs | CMR Length | p-value | Adjusted p-value | CMR $\Delta\beta$ | CHR | Location | Annotated Gene Name | Genetic Context |
| --- | --- | --- | --- | --- | --- | --- | --- | --- | --- |
| Conflict | cg27153059, cg01444541 | 2 | 3.04E-07 | 0.0139 | 0.0368 | 11 | 18719942 | *TMEM86A* | TSS1500 |
| Conflict | cg19742916, cg11753308 | 2 | 1.42E-05 | 0.149 | 0.0439 | 14 | 100086137 |  |  |
| Conflict | cg23915527, cg03441844, cg19247228 | 3 | 1.80E-05 | 0.149 | -0.1754 | 1 | 161368787 |  |  |
| Conflict | cg20011360, cg04301614 | 2 | 1.98E-05 | 0.149 | 0.0573 | 4 | 37585660 | *C4orf19* | TSS200 |
| Conflict | cg24555670, cg12174804 | 2 | 2.63E-05 | 0.149 | -0.0475 | 8 | 18244502 |  |  |
| Conflict | cg13818430, cg21839972 | 2 | 2.72E-05 | 0.149 | 0.0425 | 12 | 7904082 |  |  |
| Conflict | cg25610428, cg14792160<sup>a</sup> | 2 | 2.75E-05 | 0.149 | 0.0740 | 1 | 152758453 | *LCE1E* | TSS1500 |
| Conflict | cg11064039, cg06242242, cg05729249<sup>a</sup> | 3 | 2.92E-05 | 0.149 | -0.1740 | 7 | 766100 | *PRKAR1B*; *HEATR2* | 5'UTR; TSS1500 |
| Conflict | cg08860498, cg16308533, cg11629889, cg12559685, cg21946667, cg07871971 | 6 | 3.62E-05 | 0.166 | 0.0694 | 17 | 40838861 | *CNTNAP1* | Body |
| Conflict | cg23825301, cg11095027, cg26599989, cg14889653, cg25226711, cg26779330 | 6 | 4.53E-05 | 0.1863 | 0.1168 | 11 | 1296823 | *TOLLIP* | 3'UTR |
| Conflict | cg07969622, cg05208878, cg17336139, cg08831999, cg22871485, cg20667684, cg01966129 | 7 | 4.87E-05 | 0.1863 | 0.0717 | 22 | 22901569 | *PRAME;*<br>*LOC64869*<br>*1* | 5'UTR;<br>TSS1500 |
| Direct care | cg08344844, cg11433815, cg02313495, cg26662201 | 4 | 3.20e-06 | 0.106 | 0.0508 | 15 |  | *BMF* | Body |
| Direct care | cg05694021, cg05438336 | 2 | 6.90e-06 | 0.106 | -0.0625 | 12 |  |  |  |
| Direct care | cg02468728, cg17735531, cg06597652 | 3 | 1.31e-05 | 0.131 | -0.0519 | 19 |  | *HCN2* | Body |
| Indirect care | cg17963157, cg07452489<sup>a</sup> | 2 | 1.4e-06 | 0.0587 | -0.0624 | 16 | 23653689 | *PALB2;*<br>*DCTN5* | TSS1500<br>; Body |
| Indirect care | cg17426192, cg26550162, cg20386580 | 3 | 2.6e-06 | 0.0587 | -0.0585 | 10 | 105344747 | *NEURL* | Body |

**Note.** CMR = Co-methylated region; <sup>a</sup>mQTL

### Family dynamic-associated DNAm associated with measured and peer-ranked child health

Path analyses were conducted to simultaneously estimate the associations between identified DNAm CMRs and multiple child health outcomes. To reduce the number of variables in our path analyses, we conducted PCA on CMRs identified in the three EWASs with parental conflict, direct care, and indirect care (**Supplementary Table 2**). The top three PCs of conflict-associated CMRs accounted for 60.4% of the total variance. The loadings of CMRs on PC1 and PC2 all had consistent directionality with the EWAS associations, while that on PC3 had one opposite and one consistent directionality. The top one PC of direct care- and indirect care-associated CMRs accounted for 54.3% and 57.7%. For all CMRs loading onto these PCs, the direction of their loadings was opposite to the direction found in the EWAS associations. These five family dynamic-associated CMR PCs were included in the path models. Pearson’s correlations showed that each peer-ranked family dynamics variable was significantly correlated with their corresponding CMR PCs in the expected direction (|*r*s| = 0.44-0.74, *p*s < 0.001).

Using these five CMR PCs, we first ran a path model (**Figure 4a**) to test our second hypothesis—the differential DNAm in the identified family dynamic-associated CMRs would be linked to measured child health. The model was saturated (i.e., degree of freedom = 0) hence model fit indices were not informative. Both direct and indirect care CMR PCs were significantly associated with children’s SFT and peer-ranked child health with moderate to large effect sizes (|*B*s| = .44 – .56). Since the direct and indirect care-associated CMR PC1 were negatively correlated with their corresponding variables of interest, the negative path estimates indicated that lower levels of DNAm, reflecting higher peer-ranking of paternal caregiving, were associated with better child health (i.e., higher SFT and peer-ranked child health).

**Figure 4.**
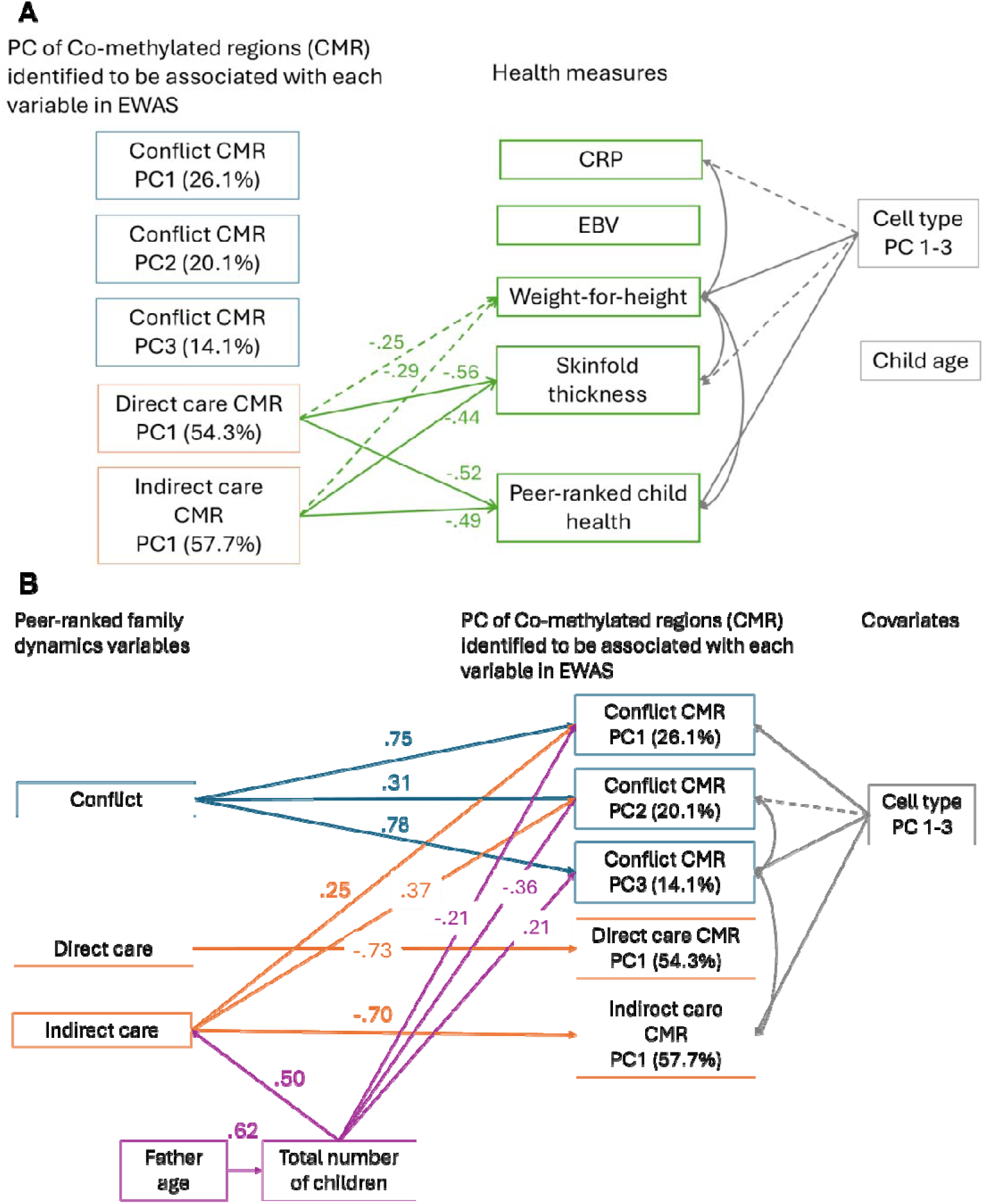
Path diagram showing the associations between the DNAm levels of identified CMR PCs with child health measures (A) and the associations between family factors and the DNAm levels of identified CMR PCs (B). In A, significant paths (p < 0.05) were indicated with solid lines and marginally significant paths (p < 0.1) were indicated with dotted lines. The green lines were paths between main variables, while the grey lines were paths with covariates. In B, the blue lines were paths from peer-ranked parental conflict; the orange lines were paths from father cares; the purple lines were paths from family factors; the grey lines were paths from covariates and covariance of residuals. Path estimates were all standardized.

### Shared contribution of conflict and indirect care on family dynamic-associated DNAm

We ran a second path model to simultaneously examine the shared contribution of all the peer-ranked family dynamics variables on identified CMRs as well as including relevant family factors (i.e., number of children and fathers’ ages) as predictors in the model (**Figure 4b; Supplementary Table 3**). The fit indices met our criteria for good fit except SRMR, χ2 (35) = 40.458, *p* = .242, CFI = .979, RMSEA = .054, and SRMR = .081. DNAm levels at conflict-associated CMR PCs were simultaneously predicted by peer-ranked parental conflict, indirect care, and total number of children. Specifically, as expected, higher peer-ranked parental conflict was linked to higher DNAm levels in all three conflict-associated CMR PCs with moderate to large effect sizes (*B*s = .31 – .78). Children of fathers who had a higher ranking in indirect care also showed higher DNAm levels in conflict-associated CMR PC1 and PC2 with small and moderate effect sizes, respectively. Children in a family with larger total number of children showed lower DNAm level of conflict-associated CMR PC1 and 2, but higher DNAm levels at PC3, again with small to moderate effect sizes (|*B*s| = .21 – .36). DNAm levels of direct and indirect care-associated CMR PCs were only explained by their corresponding peer-ranked variables, with large effect sizes (*B*s = −.73 – −.70). The path model suggested that older fathers had more children in their household, which in turn was associated with higher peer-ranking for indirect care.

## Discussion

The role of fathering in early family contexts on children’s biology and health is often understudied (Gettler, 2016; Gettler & Barr, 2022). Additionally, prior studies on the biological embedding of family dynamics have largely been conducted in Euro-American contexts in which the cultural and socioecological conditions may not be generalizable to many global populations (Gettler, Lin, et al., 2020). The current study demonstrated that differential blood DNAm in children was associated with experiences of both positive and negative family dynamics—father’s involvement and parental conflict, respectively—peer-ranked by other fathers in a smaller-scale, subsistence, Bandongo fishing-farming society in the Republic of the Congo. We found that some of these DNAm signals were linked to Bandongo children’s physical health and growth. Furthermore, parental conflict and father indirect care simultaneously contributed to conflict-associated DNAm, while paternal care-associated DNAm was not explained by other family factors beyond fathers’ care. Additionally, total number of children in the household had opposing directionality of DNAm associations as compared to parental conflict when both were considered simultaneously. Collectively, our findings suggested that paternal care and parental conflict may have comparable associations with children’s biology across socio-ecologically diverse contexts.

### Pediatric DNAm associated with family dynamics in Bandongo was linked to possible health relevance

Via a path model, we investigated the associations between CMRs identified in EWAS models with health biomarkers (CRP and EBV) and outcomes (anthropometrics and father-rated child health) also measured in this cohort. This allows us to examine the possible health relevance of the CMRs linked with parental conflict and father caregiving. Our EWAS models demonstrated that differential DNAm levels at eleven and five CMRs were associated with peer-ranked parental conflict and paternal care in the Bandongo community, respectively.

Furthermore, the paternal care-associated DNAm covaried with indicators of children’s physical growth and health. These findings extend prior work on DNAm associations with parental conflict and caregiving documented among children in the United States, Canada, and the United Kingdom (Bush et al., 2018; Chan et al., 2025; Dunn et al., 2019; Essex et al., 2013; Merrill, Moore, et al., 2021; Provenzi et al., 2020) by showing these associations can also be observed among children residing outside of industrialized, Euro-American societies, in a smaller-scale, agricultural community in an energetically-demanding environment, which represents economic, cultural, and ecological dynamics rarely explored in past studies in this area.

Based on prior ethnographic observation of the Bandongo community, parental conflict—including couples’ verbal and occasionally physical disputes—is not uncommon, yet it is still recognized to have negative impact on children (Boyette et al., 2018; Gettler, Lin, et al., 2020).

The biological toll of parental conflict on Bandongo children was first demonstrated through EAA—biological aging derived from DNAm, in which higher EAA was found in Bandongo children experiencing higher parental conflict (Gettler, Lin, et al., 2020). Our current discovery with EWAS expands on this finding and now includes paternal care-DNAm associations. Specifically, some of the CMRs identified in the current study were annotated to genes related to development and growth, partially supporting our hypotheses (H1). For example, our high-confidence CMR associated with parental conflict resided in *TMEM86A*, which encodes a protein that has been implicated in lipid metabolic process and metabolic (Cho et al., 2022). Furthermore, some identified CMRs were in genes related to stress and immunology (e.g., *TOLLIP*, which encodes a protein that regulates inflammatory signal). It is worth noting that three of the identified CMRs could be influenced by mQTLs, reflecting potential confounds between genetics and the family-level measures. While not directly comparable, this association of differential CMRs within or nearby stress-related genes is consistent with prior findings in the same cohort in which parental conflict associated with elevated EBV, a biomarker linked to stress-responsive physiology (Boyette et al., 2018). In an adult sample from urban Philippines—a low-income country with high rates of inflammation-related disease—extended parental absence, a common feature of caregiving in that context, was associated with DNAm in immune-related genes (McDade et al., 2017). Blood DNAm-family dynamics associations in genes related to stress and inflammation were also found among Canadian children (Chan et al., 2025). Together, our study provided empirical evidence of pediatric DNAm associations of both positive and negative family dynamics beyond predominantly White, Euro-American sociocultural contexts, while revealing differential DNAm patterns in genes of similar functions identified in past studies from those contexts (Chan et al., 2025; Provenzi et al., 2020).

Capitalizing on the health outcomes measured in the same cohort, we also found that DNA methylation differences in stress-related regions linked to paternal care—but not parental conflict—were associated with children’s physical growth and how peers rated their health, partly confirming our predictions (H2). Our findings align with the more recent POHaD framework that emphasizes paternal influences on children’s health and development through epigenetics (Soubry, 2018). These links can be expected given that, in the Bandongo community, fathers’ indirect care often means providing food and resources, which would naturally support children’s physical development (Boyette et al., 2018). Our findings that direct care-associated DNAm also explained variation in children’s physical growth and health even after accounting for indirect care-associated DNAm are in line with the prior findings of paternal direct care being a strong predictor of children’s health in the Bandongo community (Boyette et al., 2018). Similarly, the lack of significant associations between our conflict-associated DNAm and children’s health is consistent with prior null findings on the direct association between parental conflict and children’s energetic well-being in the same cohort (Boyette et al., 2018). While parental conflict has previously been linked to signs of elevated stress physiology in children (Boyette et al., 2018), our results suggest that DNA methylation may not mediate this relationship. Instead, it may act as an independent biomarker or operate through biological pathways beyond those we measured here.

In sum, our study provided evidence that experiencing higher levels of parental conflict and lower levels of paternal care are biologically embedded through DNAm in Bandongo children, and paternal care-associated DNAms predicted variation in slower physical growth and worse physical health, above and beyond children’s age and immune cell compositions. Although not formally tested in our study, these current findings together with previously found association between paternal care and children’s physical growth in Bandongo families (Boyette et al., 2018) make it tempting to speculate that DNAm may serve as a biological pathway linking paternal care to children’s physical growth and health.

### Family factors simultaneously explained DNAm associations with parental conflict

Taking a family system perspective, we also accounted for the family factors that may be of relevance in the specific cultural context of the Bandongo community—father’s age and total number of children—and further examined the simultaneous contributions of parental conflict, paternal caregiving, and these family factors using a path model. In this context, our hypothesis of shared contribution of parental conflict and paternal caregiving on the DNAm levels at the identified CMRs (H3) was partially supported. Specifically, DNAm levels of conflict-associated CMRs were associated with parental conflict, as expected, but also showed a significant direct effect from paternal indirect care and total number of children in the same household, with an effect of opposite directionality to parental conflict itself. These patterns suggest parental conflict-DNAm associations represent a more complex story than the DNAm associations with paternal care and warrants a closer inspection. Given the independent kin network in the Bandongo community, larger families with more children are generally valued since they can contribute to subsistence and household labor, including older children caring for their siblings. Given the importance of sibling care and relationships across development in Bandongo communities, sibling support can also play a significant role in the family system, as in other contexts (Feinberg et al., 2012). Prior studies on sibling relationships in White, Euro-American samples have shown that sibling relationships can buffer the negative effect of interparental conflict, potentially through providing a source of security, support, and companionship during the stressful experience (Davies et al., 2019). To date, however, research focused specifically on sibling buffering of parental conflict remains understudied outside of Euro-American contexts (Feinberg et al., 2012). In our path model, we illustrated that while parental conflict increased the DNAm levels of the identified CMRs, the number of siblings have an opposing effect on most of these DNAm levels (except PC3, which is less interpretable as explained earlier), above and beyond parental conflict itself. Putting this together, children who experienced higher level of parental conflict showed higher level of DNAm at the identified CMRs, while those who have more siblings showed a lower level of DNAm than their counterparts who have fewer siblings, when both factors were considered simultaneously. Although a moderating effect was not formally tested in our model due to limited statistical power, it is plausible that the number of siblings could buffer the effect of parental conflict on DNAm.

Paternal care-associated CMRs were uniquely associated with the corresponding peer-ranked care variable, without any other family variables in our model explaining additional variances. Although we found that father’s age was related to total number of children, which in turn linked to peer-ranked paternal indirect care (i.e., fathers being ranked as working harder), we did not find a direct effect from father age to any of the peer-ranked variables. Nonetheless, this finding supports peer rankings in this study was not driven by the social status of the fathers, which is often linked with age, and lends support to the reliability of this culturally appropriate measure. Interestingly, peer-ranked indirect care was also positively linked to differential DNAm levels of conflict-associated CMRs, above and beyond parental conflict (same direction) and total number of children (opposite direction). These associations seem counterintuitive and our current study does not provide a straightforward explanation. In Bandongo society, some forms of fathers’ indirect caregiving involve competition for status with other men and forms of physical risk taking (e.g., climbing tall trees to extract resources) (Boyette et al., 2018; Gettler et al., 2019; Gettler, Lew-Levy, et al., 2020). Although it is speculative, it is possible that children with fathers who were oriented towards competition or risk-taking experienced family dynamics that helped shape the observed patterns for DNAm at conflict-associated CMRs.

## Limitations

There are a few limitations of this study that should be considered. First, our results were based on cross-sectional data, therefore, the associations observed in our EWAS models and path models were correlational in nature. There was no temporal precedence in our predictors and outcome variables. The directionality of the associations in our path model was specified based on the theories of biological embedding. However, it is possible that those DNAm patterns could exist before the exposure of the measured family dynamics or the two could have shared causes that were not captured in the model. Longitudinal data will be needed in future studies to ascertain that the DNAm patterns associated with family dynamics can be attributed to the exposure of family dynamics by controlling for earlier DNAm profiles. Additionally, it is important to note that DNAm in peripheral tissues often function as biomarkers of environmental exposure and physiological state rather than direct regulatory mechanisms. The associations observed here may reflect biological correlates of family dynamics, such as differences in stress physiology or immune activity, rather than causal molecular pathways linking family experience to health. Our findings are also specific to blood samples, which primarily consist of immune cells. Second, our study in the Bandongo community had a smaller sample size compared to some recent EWASs on family environment using large-scale, Euro-American cohorts. However, this reflects the realities of working in smaller-scale subsistence societies, where communities have low population sizes, that have largely been absent to-date from developmental research focusing on biological embedding of social experiences, including in family life. We accounted for family interdependence (17 families for the 54 children) using robust lme models given that our key predictors were measured at the household level and shared by children in the same household. This limited our ability to separate between-family and within-family (child-level) effects. Because paternal peer ratings were available for 17 families, analyses of key predictors should be interpreted as most sensitive to larger associations, with limited ability to detect small but potentially meaningful effects. In spite of that, our findings of DNAm associations with parental conflict and father’s caregiving aligned with the current literature of family adversity and maternal caregiving, predominantly in White, Euro-American samples (Chan et al., 2025; Provenzi et al., 2020) and provide empirical evidence that the biological embedding of family factors shows parallels across socio-ecologically diverse contexts. Lastly, the ranking measures in our study did not differentiate the types of conflicts/care as well as the severity of conflicts and our study did not include maternal reports of family dynamics. Future studies taking a family system approach should investigate the roles of both mothers and fathers and other caregivers in children’s biology and health.

## Conclusions

The current study provides evidence of the biological embedding of parental conflict and father caregiving through DNAm in a small, fisher-farmer subsistence community in the Republic of the Congo, demonstrating cross-cultural commonalities in this biological process. Importantly, our results highlighted the importance of considering other family factors, such as the presence of siblings and supported the implications of these DNAm associations for children’s health and development. Together, these findings underscore the importance of examining family relationships within diverse socioecological contexts to better understand how early social experiences become biologically embedded.

## Author Note

We offer our immense gratitude to the Bandongo families who made this study possible as well as the leaders of the local community, the Precôt and Secrétaire. We are similarly grateful to those in Congo who helped facilitate this research: The Institut National de Recherche en Sciences Exactes et Naturelles (IRSEN), especially Dr. Clobite Bouka Biona; the Centre de Recherche et D’Etudes en Sciences Sociales et Humaines (CRESSH), especially Dr. Francois Ibara; Dzabatou Moise and Mindoula Koutain for their assistance with fieldwork.

The authors of this study have no affiliations with nor involvement in any organization or entity with any financial interests or non-financial interests in the subject matter or materials discussed in this manuscript.

This work was supported by the Jacobs Foundation, Welcome Trust.

## Notes

### Competing Interest Statement

The authors have declared no competing interest.

