## Supplementary Material for "Children’s DNA Methylation and Family Dynamics in a Congo Basin Subsistence Community: Links with Parental Conflict and Fathers’ Caregiving"

**Supplementary Table 1.** Descriptive statistics of demographic, family dynamics, and child health variables

**Supplementary Table 2**. Loadings of the CMRs identified to be associated with each family dynamic variable on their top PCs

**Supplementary Table 3.** Full estimates for path model 1 and 2 (separate excel sheet)

**Supplementary Figure 1.** Estimated proportions of six cell types in blood: neutrophils, monocytes, B lymphocytes (Bcells), T helper lymphocytes (CD4T), T cytotoxic lymphocytes (CD8T), and natural killer lymphocytes (NK).

**Supplementary Figure 2.** Variable co-methylated region (CMR) length distribution (A) and CMR β value distribution (B) of the 54 blood samples.

**Supplementary Figure 3.** Pearson’s correlation heatmap of all main variables**.**

**Supplementary Figure 4.** Genetic control of identified CMRs (mQTL)

**Supplementary Figure 5.** Full, hypothetical path model testing the simultaneous associations between peer-ranked variables, DNA methylation at identified co-methylated regions, and family factors.

**Supplementary Method.** Detailed methods for DNA methylation preprocessing, cell type estimation, and construction of co-methylated regions

**Supplementary Table 1.** Descriptive statistics of demographic, family dynamics, and child health variables

|  | Mean | SD | Range (min-max) | N |
| --- | --- | --- | --- | --- |
| Age (years) | 8.48 | 3.83 | 2-17 | 54 |
| Sex (female ratio %) | 51.85 | - | - | 54 |
| Father's age at sampling (years) | 38.29 | 7.66 | 24-50 | 17 |
| Father's age on children's birth year (years) | 31.65 | 6.68 | 13-46 | 54 |
| Mother's age at sampling (years) | 34.24 | 6.65 | 22-48 | 17 |
| Mother's age on children's birth year (years) | 26.98 | 6.48 | 11-42 | 52 |
| Total number of children per household | 6.29 | 3.44 | 2-15 | 17 |
| Parental conflict ranking | 3.18 | 1.23 | 1.12-5.38 | 17 |
| Father as provider ranking | 8.63 | 3.64 | 1.6-15.38 | 17 |
| Father as caregiver ranking | 5.2 | 2 | 1.39-8.42 | 17 |
| Children's health ranking | 6.15 | 2.13 | 2.19-9.5 | 17 |
| Weight-for-height (z-score) | 0 | 1 | -1.57-2.7 | 54 |
| triceps skinfold thickness (z-score) | 0 | 1 | -2.19-3.01 | 54 |
| EBV antibody titers (AU/ml) | 192.21 | 202.87 | 24.62-693.52 | 48 |
| C-reactive protein level (mg/L) | 2.94 | 4.06 | 0-16.34 | 52 |

**Supplementary Table 2**. Loadings of the CMRs identified to be associated with each family dynamic variable on their top PCs

| **Conflict-associated CMRs** | **PC1** | **PC2** | **PC3** |
| --- | --- | --- | --- |
| cg25610428 | 0.76^a^ | -- | -- |
| cg23825301 | -- | 0.83^a^ | -- |
| cg27153059 | -- | 0.55^a^ | -- |
| cg13818430 | 0.59^a^ | -- | -- |
| cg19742916 | -- | 0.61^a^ | -0.71^b^ |
| cg08860498 | -- | 0.73^a^ | -- |
| cg07969622 | 0.58^a^ | -- | -- |
| cg20011360 | -- | -- | 0.82^a^ |
| cg11064039 | -0.91^a^ | -- | -- |
| cg24555670 | -0.64^a^ | -- | -- |
| cg23915527 | -- | -- | -- |
| **Direct care-associated CMRs** | **PC1** |  |  |
| cg05694021 | 0.438^b^ | -- | -- |
| cg08344844 | -0.845^b^ | -- | -- |
| cg02468728 | 0.851^b^ | -- | -- |
| **Indirect care-associated CMRs** | **PC1** |  |  |
| cg17426192 | 0.76^b^ | -- | -- |
| cg17963157 | 0.76^b^ | -- | -- |

*Note*. Only loadings larger than 0.4 are presented. Only one CpG was listed for each CMR.
^a^ Same directionality between loading and association with variable of interest in the EWAS conducted in the current study; ^b^ Opposite directionality between loading and association with the variable of interest in the EWAS conducted in the current study

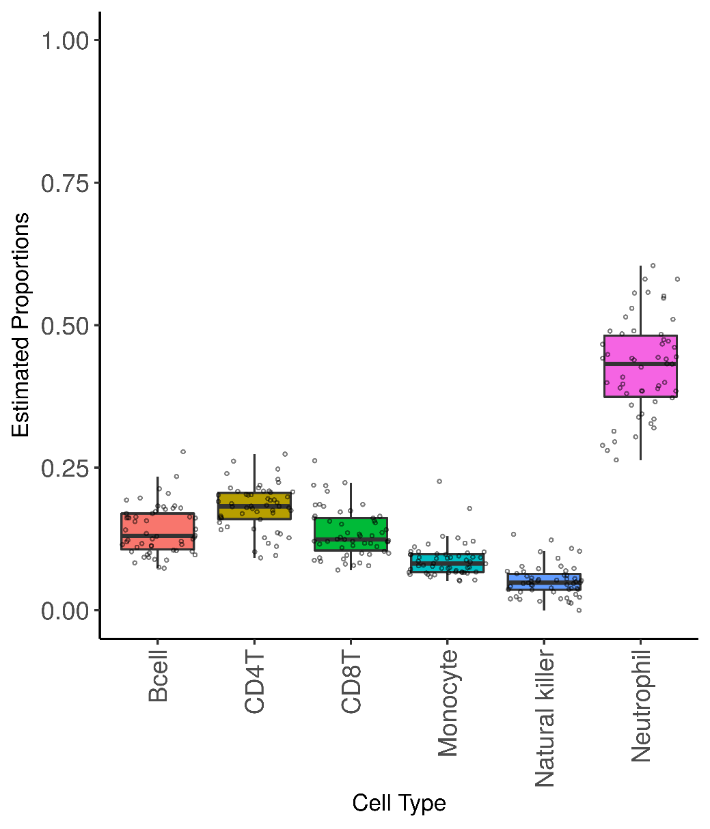

**Supplementary Figure 1.** Estimated proportions of six cell types in blood: neutrophils, monocytes, B lymphocytes (Bcells), T helper lymphocytes (CD4T), T cytotoxic lymphocytes (CD8T), and natural killer lymphocytes (NK). Relative cell type proportions of each sample were predicted by the constrained projection method (Houseman et al., 2012), with IDOL reference library (Salas et al., 2018) by using the estimateCellCounts2 function in the R package FlowSorted.Blood.EPIC (version 2.2.0).

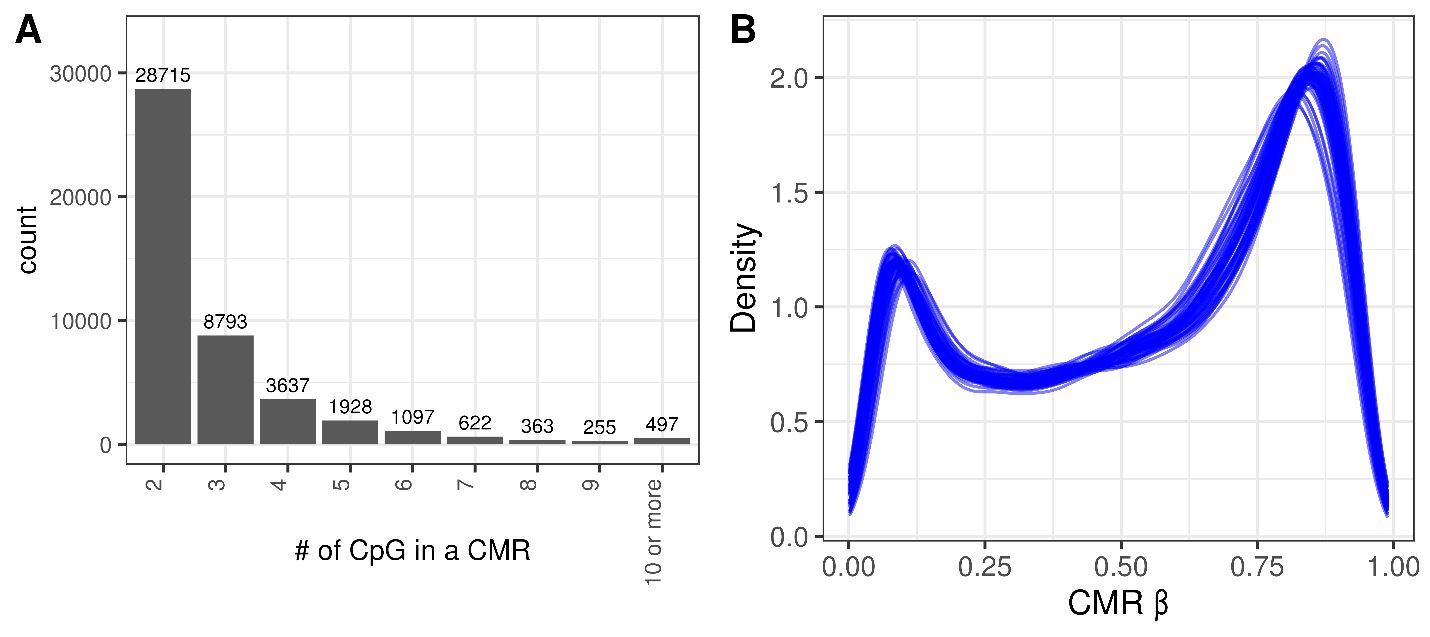
**Supplementary figure 2.** Variable co-methylated region (CMR) length distribution (A) and CMR β value distribution (B) of the 54 blood samples.

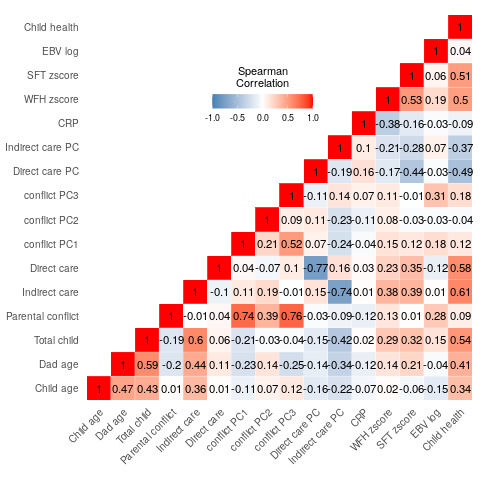

**Supplementary Figure 3.** Pearson’s correlation heatmap of all main variables**.** WFH = weight-for-height; SFT = skinfold thickness; EBV = Epstein‐Barr virus; CRP = C‐reactive protein levels. Pearson’s correlations showed that the peer-ranked variables were not significantly correlated with each other (*r*s = -0.09 – 0.02, *p*s > 0.05). Child health and SFT were positively correlated with both direct and indirect father cares (*r*s = 0.35-0.63, *p*s < 0.01), while WFH was also positively correlated with indirect care (*r* = 0.4, *p* = 0.003). Furthermore, father’s age was significantly correlated with number of children in the household (r = 0.62, p < 0.0001), and both father’s age and number of children were correlated with peer-ranked indirect father care (*r*s = 0.61 and 0.49, *p*s = 0.0001 and <0.0001, respectively). Direct and indirect care CMR PCs (higher levels in these PCs were linked to lower peer-ranked care) were negatively correlated with SFT and peer-ranked child health (*r*s = -0.42 - -0.29, *p*s = 0.001 – 0.03).

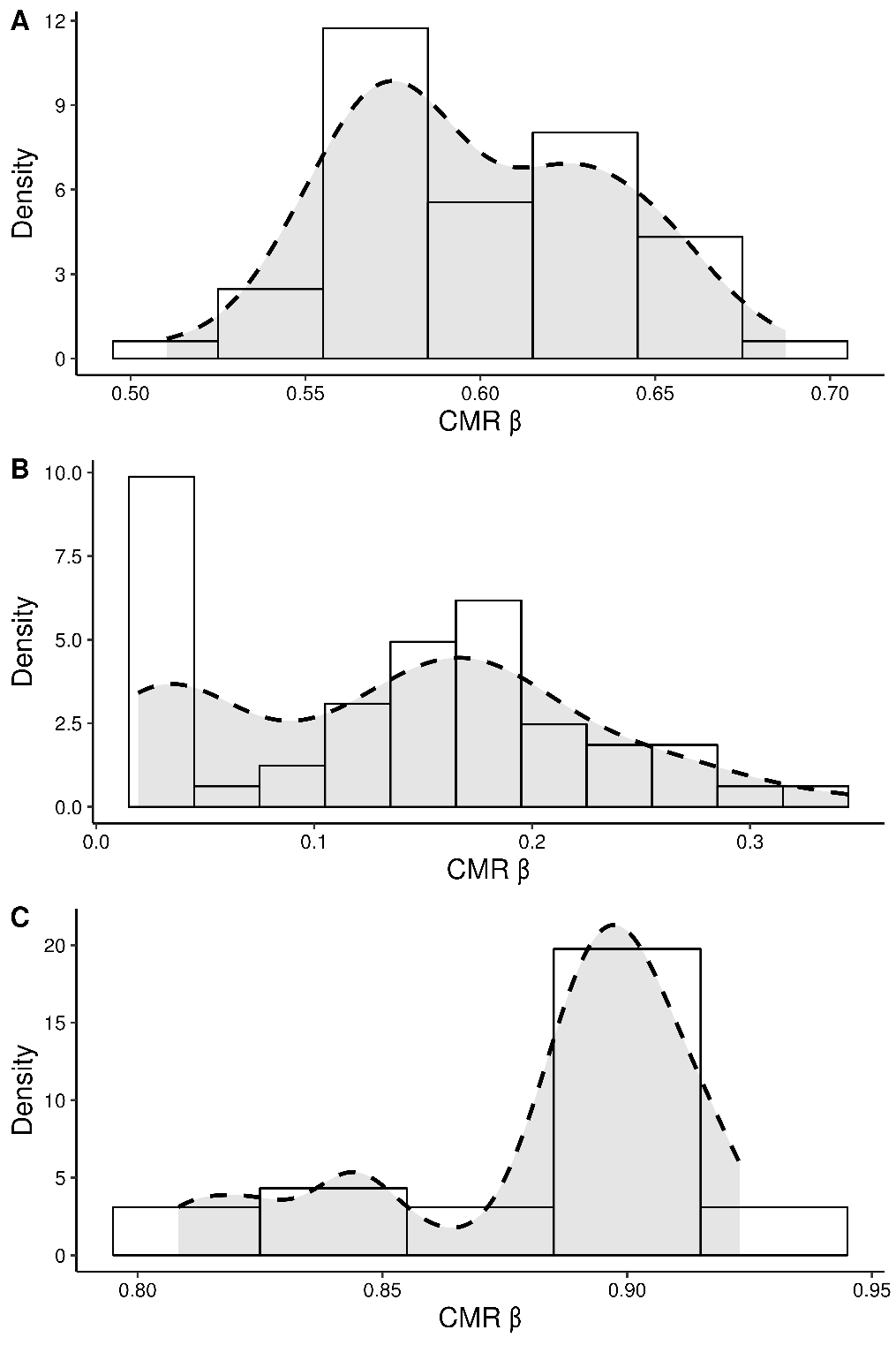

**Supplementary Figure 4. Genetic control of identified CMRs (mQTL)** (A) CMR cg25610428 and cg14792160 (B) CMR cg11064039, cg06242242, cg05729249 (*PRKAR1B; HEATR2*) and (C) CMR cg25610428, cg14792160 have bimodal CMR β distribution.

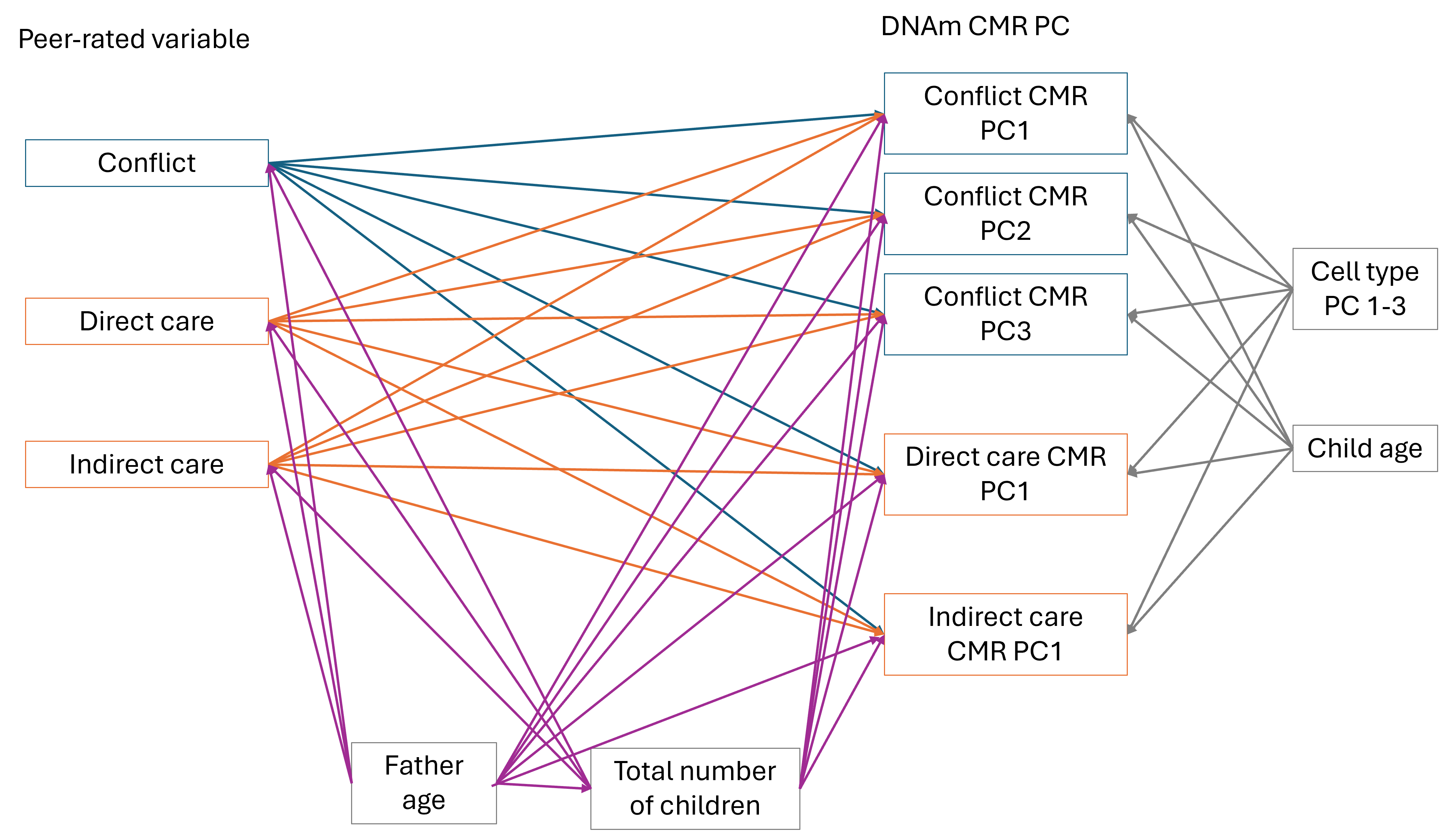

**Supplementary Figure 5.** Full, hypothetical path model testing the simultaneous associations between peer-ranked variables, DNA methylation at identified co-methylated regions, and family factors. The blue lines were paths from peer-ranked parental conflict; the orange lines were paths from father cares; the purple lines were paths from family factors; the grey lines were paths from covariates. All residuals of the CMR PCs were allowed to covary with each other but were not shown on the figure for readability.

**Supplementary Method**

**Co-methylated Regions as Unit of Analysis**

Due to the ultra-high dimensional nature of probe-wise DNAm data coupled with the inherent limitations on sample size in unique populations, including our samples in which we have recruited all eligible participants from a small village, potential biological signals can be missed given the heavy multiple test correction burden. Region-based analysis reduces redundant measurements and noise while capturing patterns that are consistent and more likely to be biologically relevant in the data (Gatev et al., 2020). Closely located cytosine-guanine (CpG) dinucleotides sites tend to have correlated methylation states and are more likely functioned as biological units (Gatev et al., 2020; Hui et al., 2018). Thus, in the current study, the unit of analysis employed was co-methylated regions (CMRs), which are defined as groups of adjacent CpGs with correlated DNAm methylation levels across samples (Gatev et al., 2020). CMRs were constructed using the cmr function in the CoMeBack R package (version 1.0) (Gatev et al., 2020). Adjacent two or more CpG array probes were grouped together as a CMR if they met both of the following criteria: 1. probes were within 1000 base pairs (bp) distance and were chained by unmeasured CpGs in the genomic background within 400bp; 2. DNAm β value of adjacent probes across samples has a Pearson correlation coefficient of 0.4 or higher. Following these steps, a total of 50,215 CMRs were constructed from 744,516 probes in this cohort. A composite DNAm β value of a CMR (CMR β) was represented by the median DNAm β value of all CpGs within the CMR. To meet the equal likelihood assumption of Benjamini-Hochberg multiple test correction (Korthauer et al., 2019), an interquantile range filter was applied to select CMRs with CMR β value with ≥ 2.5% variation across samples in the 10^th^ and 90^th^ percentile. A final count of 45,907 variable CMRs was included in the association analysis.

***Children’s biomarkers***

As previously described in detail (see Boyette et al. and Gettler et al. 2021), blood by finger-prick sampling was stored as dried blood spots (DBS) on filter paper. DBS samples were assayed for EBV and CRP levels using enzyme-linked immunosorbent assay approaches at the Global Health Biomarker Laboratory at the University of Oregon. We eliminated one individual based on an EBV value that was 6+ SD above the sample mean. One individual was EBV seronegative. Intra- and inter-assay coefficients of variation for the CRP ELISAs were 3.6% and 6.3%, respectively; for the EBV IgG ELISAs, they were 7.5% and 13.7%, respectively. EBV antibody levels were strongly right-skewed and log-transformation was applied to reduce skewness. Samples that had CRP levels > 3 mg/L were regarded as having elevated CRP level, which indicated the presence of chronic low-grade inflammation low‐grade immune activation and active acute infection (see Boyette et al. 2018 and Gettler et al. 2021).
